# Pathologic profibrotic activation of distinct myeloid cell subsets in a model of impaired healing after anterior cruciate ligament reconstruction

**DOI:** 10.1101/2025.07.31.667715

**Authors:** Wataru Morita, Yuki Suzuki, Ruoxi Yuan, Scott A. Rodeo, Kyung-Hyun Park-Min, Lionel B. Ivashkiv

## Abstract

Anterior cruciate ligament reconstruction (ACLR) may fail due to poor bone-to-tendon integration, yet the cellular mechanisms underlying this process remain incompletely understood. Using a murine model of ACLR, we show that host-derived immune cells are the primary drivers of early healing, while tendon graft-derived stromal cells also contribute at the tendon-bone interface. We developed a model of impaired post-ACLR healing by a surgical approach, that led to increased infiltration of immune cells at the tendon-bone interface, particularly CX3CR1+ monocytes/macrophages (MonoMacs). Single-cell RNA sequencing revealed dynamic monocyte-to-macrophage transitions, with an enrichment of fibrotic, mechanosensitive CX3CR1+ MonoMacs in the impaired healing model. These findings suggest that excessive recruitment or functional shifts in host-derived MonoMacs may disrupt normal healing dynamics. Our study identifies CX3CR1+ MonoMacs as a key cell population related to inferior bone-to-tendon healing after ACLR and potential therapeutic targets to improve surgical outcomes.

## INTRODUCTION

Anterior cruciate ligament (ACL) injuries make up nearly a half of all knee injuries, and are becoming more frequent, particularly among young adults involved in high-intensity sports^1^. While physiotherapy may be the initial approach especially for recreational athletes^2^, full recovery is often unachievable with conservative treatment or primary repair, which has a failure rate exceeding 40 %^3^. As a result, anatomical reconstruction using tendon autografts or allografts placed through bone tunnels in the femur and tibia is now the standard of care^2^.

Following transplantation, the tendon graft undergoes sequential healing phases, including formation of a fibrovascular membrane between the tendon graft and surrounding bone tissue, appositional bone growth to integrate with the graft, and remodeling of the tendon graft, a process known as ligamentization^4–7^. ACL reconstruction (ACLR) fails in up to 12% of patients, which can be partially due to insufficient bone-to-tendon integration or to poor remodeling of the intraarticular portion of the graft^8, 9^. Surgical technique is a major contributor of ACLR^9^. Clinical studies have reported higher failure rates when using hamstring grafts of smaller diameters^10–12^. Grafts that are relatively smaller than the bone tunnel diameter can lead to micromotion within the tunnels, a phenomenon commonly referred to as the graft piston or “windshield wiper” effect^13, 14^. Furthermore, ACL injuries significantly increase the long-term risk of post-traumatic osteoarthritis (PTOA) regardless of the treatment^15, 16^. The rate of PTOA after ACLR is approximately 50% at 15-20 years^17^. Thus, there is an unmet medical need to improve post-ACLR healing and function to decrease graft failure and PTOA development.

ACLR failure remains a significant clinical challenge, and ongoing research continues to explore the mechanisms underlying ACLR healing. Previous studies have demonstrated that host-derived cells contribute to the bone-to-tendon healing within the bone tunnels, but the infiltrating cell types were not specified^18, 19^. Moreover, many studies have used decellularized allografts and the potential contribution of tendon derived cells has not been fully addressed^20–24^. Both immune and stromal cells such as mesenchymal progenitors have been identified in tendon-bone interface tissue, and can interact in a complex and dynamic manner^25–33^. Our previous work highlighted the important role of macrophages in bone-to-tendon healing following ACLR^34, 35^. We demonstrated that depleting myeloid cells using clodronate liposomes enhanced bone-to-tendon integration, as evidenced by improved biomechanical properties and increased bone formation within the bone tunnels^34^. Furthermore, we characterized the distinct monocyte/macrophage populations in a mouse ACLR model, and identified a key population expressing chemokine receptors CX3CR1 and CCR2 that gradually accumulate after surgery and may negatively contribute to ACLR healing^35^.

In this study, we used flow cytometry and single cell RNA sequencing to define the monocyte/macrophage populations that accumulate within the healing bone tunnels after ACLR, and developed a new model of impaired post-ACLR healing that resulted in PTOA. Our study showed increased infiltration of host-derived immune cells, particularly CX3CR1-hi monocyte/macrophage cells that likely correspond to non-classical monocytes, was associated with impaired post-ACLR healing. Monocyte/macrophage populations within the bone tunnels expressed elevated inflammatory and IFN signatures^35^; impaired post-ACLR healing was also associated with increased expression of TGF-β pathway genes. These results suggest that targeting monocyte/macrophage populations may represent a promising strategy to enhance healing after ACLR surgery, and targeting non-classical monocytes may be beneficial in the setting of poor post-ACLR healing.

## RESULTS

### Host-derived immune and tendon graft-derived stromal cells are predominant cell types in bone tunnels after ACLR

To investigate the host versus graft contribution of cells in healing post-ACLR tissues, we employed a mouse model in which a tendon graft harvested from a MIP-GFP mouse (which expresses GFP only in pancreas) was transplanted into a UBC-GFP host that expresses GFP ubiquitously in all cells (Fig. 1a). Bone tunnel tissues were harvested immediately after (day 0) and 1, 2, 4 and 8 weeks post-operatively and analyzed by histology and flow cytometry; in this system all green fluorescent cells originate from the host.

**Figure 1.**
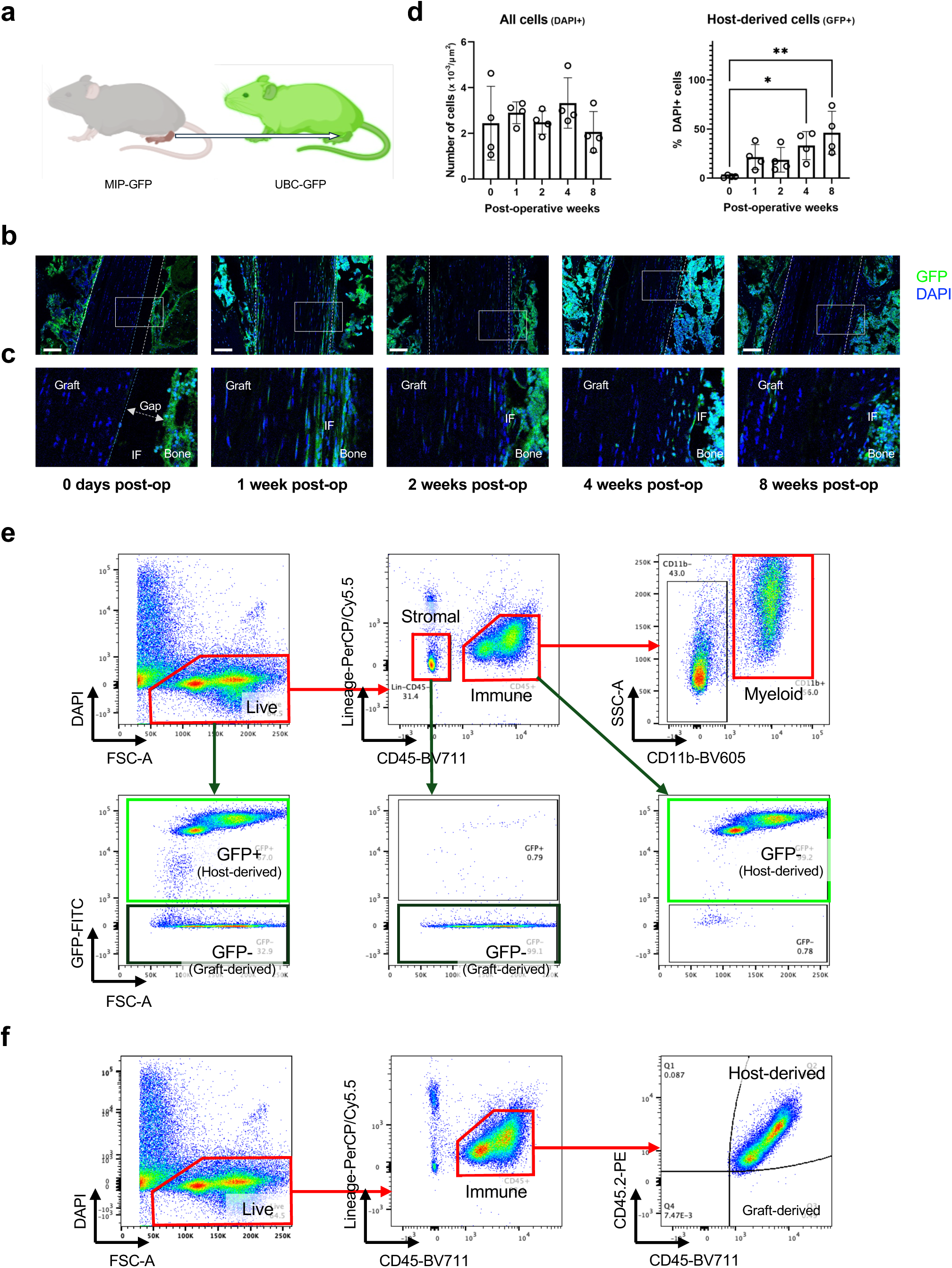
Host-derived immune cells are the predominant cells in bone tunnels after ACLR. **a**, Schematic representation of the MIP-GFP tendon graft/UBC-GFP host transplant model. **b**, Representative immunofluorescence images from 4 replicates using a MIP-GFP tendon graft/UBC-GFP host transplant model. Dotted white lines indicate the bone tunnel. Dotted blue lines indicate the edge of the tendon graft. Scale bar, 100 μm. **c**, Enlarged view of the outlined region (white box) in **b**. **d**, Quantification of number of immunofluorescent cells (n = 4 per timepoint) show no changes in total number of cells (DAPI+ cells), but gradual increase of host-derived cells (GFP+ cells) within the bone tunnels post-operatively. **e**, Representative flow cytometric plots from 3 replicates at 2 weeks post-operatively using a MIP-GFP tendon graft/UBC-GFP host transplant model showing that majority of the cells within the bone tunnels are derived from the host. Immune cells that are mostly myeloid cells are predominantly host-derived, whilst lineage (Ter119, CD31, CD3, B220, NK1.1)- CD45-stromal cells are mainly tendon graft-derived. **f**, Representative flow cytometric plots from 3 replicates at 2 weeks post-operatively using a CD45.1+ Pep Boy tendon graft/ CD45.2+ WT host transplant model showing that immune cells within the bone tunnels are predominantly derived from the host. Bars in the graphs represent the mean and standard deviation. \**p* < 0.05, \*\**p* < 0.01 by one-way ANOVA with Tukey’s post hoc test. GFP, green fluorescent protein; DAPI, 4’,6-diamidino-2-phenylindole; IF, tendon-bone interface; Graft, tendon graft; FSC, forward scatter; SSC, side scatter.

Histological analysis revealed progressive narrowing of the bone tunnel, reflecting formation of fibrovascular interface tissue and ingrowth of bone, in accordance with previous work^26, 34, 36–46^. At 1 week post-operatively, host-derived GFP+ cells were primarily localized to the tendon-bone interface, suggestive of migration from adjacent bone and bone marrow (Fig. 1b, c). The host cells persisted at the 2, 4 and 8 week post-operative time points; quantitative analysis revealed no significant change in total cell numbers within the bone tunnels over time, but a gradual increase in the proportion of host-derived cells, reaching statistical significance at 4 weeks post-operatively (Fig. 1d). To further characterize the host-derived cells we performed flow cytometry analysis on cells extracted from tissues harvested from the bone tunnels, including the tendon graft and the tendon-bone interface, at the 2-week time point. The majority of the cells present were GFP+ and thus host-derived (Fig. 1e, lower left panel). Bone tunnel cells were then separated into immune and stromal cells based upon expression of CD45 (Fig. 1e, middle panel). CD45+ immune cells were almost entirely GFP+ and thus host-derived (Fig. 1e, lower right panel), and the majority expressed the myeloid lineage marker CD11b (Fig. 1e, upper right panel). These results were the corroborated using transplants of CD45.1+ tendon grafts from Pep Boy mice into CD45.2+ hosts (Fig. 1f). In contrast to immune cells, CD45-stromal cells were largely tendon graft-derived (Fig. 1e, lower middle panel). Collectively, the results suggest that host immune cells infiltrate interface tissues post ACLR, where they can interact with graft stromal cells.

### Tendon graft-derived stromal cells accumulate at the tendon-bone interface

The current paradigm is that after ACLR tendon graft-derived cells are gradually lost over time, while host-derived cells populate the ligamentizing graft^18–24, 47^. However, there is limited knowledge about the source and role of different stromal cells at earlier time points after ACLR. Our flow cytometry results showing a dearth of host-derived and abundance of tendon graft-derived stromal cells in bone tunnels prompted us to focus on the tendon graft-derived cells in the 1-8 weeks post-operative timeframe. We employed a reciprocal model in which a tendon graft harvested from a UBC-GFP mouse was transplanted into a MIP-GFP host, and examined the bone tunnels, especially the tendon-bone interface (Fig. 2a). Notably, at the 1 week post-operative time point tendon graft-derived GFP+ cells were apparent not only in the tendon body, but showed the strongest fluorescence signal at the tendon-bone interface (Fig 2b, c). GFP+ cells persisted at the tendon-bone interface for up to 8 weeks; quantitation of GFP+ cells in 100 μm regions of interest spanning the tendon-bone interface along the bone tunnel showed a trend towards increasing cell numbers at the 1 and 2 week time points (Fig. 2d). GFP+ cells also increased in the tendon body, which is consistent with cell activation.

**Figure 2.**
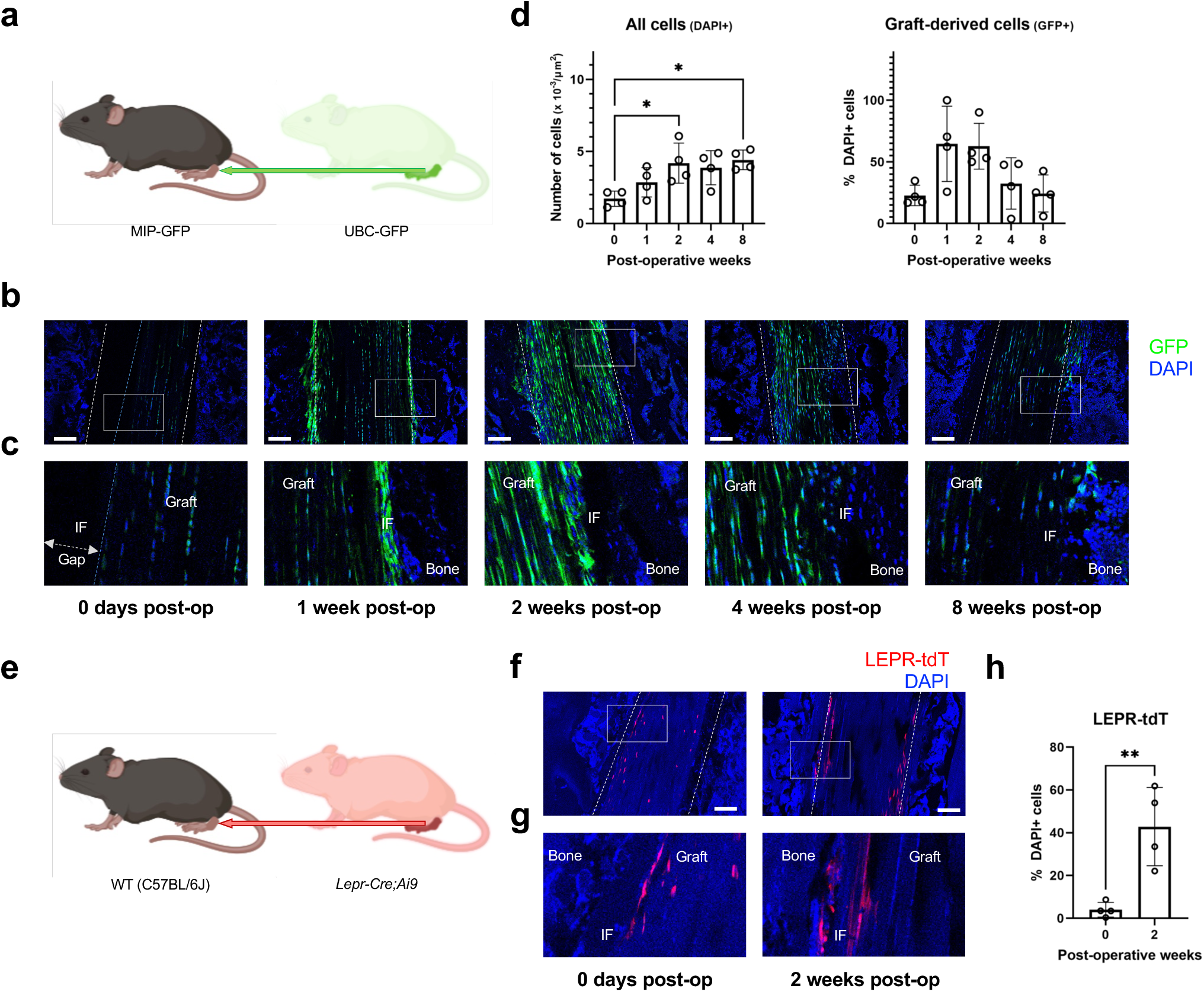
Tendon graft-derived stromal cells accumulate at the tendon-bone interface after ACLR. **a**, Schematic representation of the UBC-GFP tendon graft/MIP-GFP host transplant model. **b**, Representative immunofluorescence images from 4 replicates using a UBC-GFP tendon graft/MIP-GFP host transplant model. Dotted white lines indicate the bone tunnel. Dotted blue lines indicate the edge of the tendon graft. Scale bar, 100 μm. **c**, Enlarged view of the outlined region (white box) in **b**. **d**, Quantification of immunofluorescent cells (n = 4 per timepoint) suggesting gradual increase of cells (DAPI+ cells) at the tendon-bone interface at the early phase of healing (up to 2 weeks post-operatively). Tendon graft-derived cells (GFP+ cells) appear to increase at the tendon-bone interface at the early phase of healing (until 2 weeks post-operatively). **e**, Schematic representation of the *Lepr-Cre;Ai9* tendon graft/WT host transplant model. **f**, Representative fluorescence images from 4 replicates using a *Lepr-Cre;Ai9* tendon graft/WT host transplant model. Dotted lines indicate the bone tunnel. Scale bar, 100 μm. **g**, Enlarged view of the outlined region (white box) in **f**. **h**, Quantification of fluorescence (n = 4 per timepoint) shows increased tendon graft-derived cells (LEPR-tdT+ cells) at the tendon-bone interface at 2 weeks post-operatively compared to immediately after ACLR surgery. Bars in the graphs represent the mean and standard deviation. \**p* < 0.05 by unpaired *t* test. IF, tendon-bone interface; Graft, tendon graft; DAPI, 4’,6-diamidino-2-phenylindole.

To validate this finding, we used another transplant model in which a tendon graft harvested from a *LepR-Cre;Ai9* mouse was transplanted into a WT host (Fig. 2e). *Lepr* is expressed in a subset of ligament- and tendon-derived cells including fibroblasts (Supplementary Fig. 1). Consistent with the UBC-GFP tendon graft/MIP-GFP host transplant model, we observed tendon graft-derived cells at the tendon-bone interface at 2 weeks post-operatively (Fig. 2f, g). Image quantification confirmed a significant increase in tendon graft-derived cells at the tendon-bone interface compared to baseline (Fig. 2h). Furthermore, TUNEL staining showed minimal cell death of tendon graft-derived cells in the bone tunnels beyond the immediate post-operative period, indicating that these cells do not rapidly disappear following ACLR (Supplementary Fig. 2).

Collectively, these results suggest that tendon graft-derived cells may contribute to ACLR healing during the 8 week post-operative period. They are also consistent with our study reporting that decellularization of the tendon graft, which would result in a healing process driven predominantly by host-derived cells, significantly impairs healing outcomes (manuscript submitted for publication).

### Model of impaired post-ACLR healing leading to PTOA

To elucidate key mechanisms associated with poor ACLR outcomes, we developed a clinically relevant model of impaired post-ACLR healing, where poor integration of the tendon graft in the bone tunnel predisposes to graft failure and PTOA. This model involves creating larger bone tunnels and applying no pretension prior to graft fixation (Fig. 3a). Mechanical testing at 4 weeks post-operatively revealed significantly lower ultimate load to failure in this model, confirming inferior mechanical integrity and thus impaired post-ACLR healing (Fig. 3b). Gait analysis served as an analog to clinical outcomes^48^, and showed no significant differences in the major parameters including stride, stride length, or stride frequency between the two groups at this timepoint, suggesting that the impaired healing remained subclinical, similar to the typical clinical scenario (Fig. 3c). However, plain radiographs at 4 weeks post-operatively showed accelerated PTOA development in the impaired healing model compared to standard ACLR, particularly evident through narrowing of the joint spaces (Fig. 3d, e).

**Figure 3.**
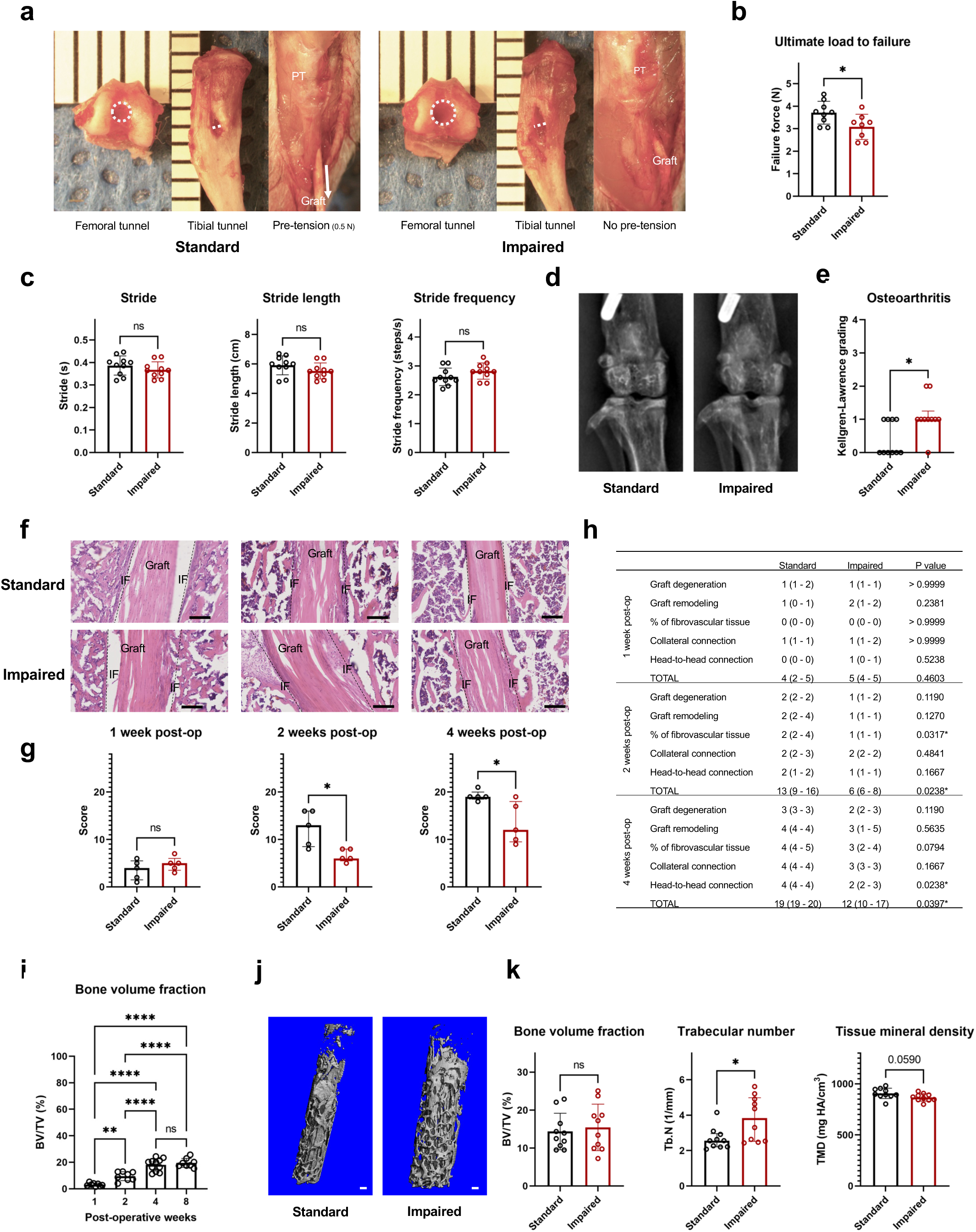
Model of impaired post-ACLR healing. **a**, Surgical approach of the standard ACLR and the impaired healing model: larger bone tunnels were created for both the femoral and tibial bone tunnels, and no pretension was applied prior to graft fixation in the impaired healing model (impaired). Dotted circles indicate the femoral tunnel intra-articular (distal) apertures. Dotted white lines indicate the tibial tunnel extra-articular (distal) apertures. The white arrow indicates that pre-tension was applied in standard ACLR. **b**, Biomechanical testing shows significantly decreased ultimate load to failure in the impaired healing model (n = 8) compared to standard ACLR (n = 9) at 4 weeks post-operatively. **c**, Gait analysis (n = 10 per group) shows no significant differences between the two models at 4 weeks post-operatively. **d**, Representative anteroposterior view plain radiographs out of 10 replicates of knee joints in the two models at 4 weeks post-operatively. **e**, Kellgren-Lawrence grading (n = 10 per group) shows higher osteoarthritis development in the impaired healing model. **f**, Representative Haematoxylin and eosin (H&E) staining out of 5 replicates of the tibial bone tunnel in the two models at 1, 2 and 4 weeks post-operatively. Dotted black lines indicate the bone tunnel. Scale bar, 200 μm. **g**, Tendon-bone tunnel healing scores of H&E-stained tissue sections (n = 5 per group) show abnormal healing in the impaired healing model from 2 weeks post-operatively. **h**, Individual scores for each parameter of the scoring system show that the inferior healing is attributed to increased fibrovascular tissues from 2 weeks post-operatively. Data shown as the median and interquartile range. **i**, μCT analysis (n = 10 - 12 per group) on new bone formation measured by bone volume fraction (BV/TV) in the tibial tunnel shows increase until 4 weeks post-operatively. **j**, Representative 3D-reconstructed μCT of newly formed bone in the tibial tunnels in the two models at 4 weeks post-operatively. Scale bar, 100 μm. **k**, μCT analysis (n = 10 per group) shows comparable BV/TV but increased trabecular number (Tb.N) in the impaired healing model. Bars in the graphs represent the mean and standard deviation (**b**, **c**, **i**, **k**) or the median and interquartile range (**e**, **g**). \**p* < 0.05; \*\**p* < 0.01; \*\*\**p* < 0.001; \*\*\*\**p* < 0.0001; ns, no significance by unpaired *t*-test (**b**, **c**, **k**), Mann-Whitney *U*-test (**e**, **g**, **h**), or one-way ANOVA with Tukey’s post hoc test (**i**). PT, patellar tendon; Graft, tendon graft; IF, tendon-bone interface.

Since anterior tibial translation may increase with compromised ACL function^49^, we also examined unstressed lateral view plain radiographs. No significant differences were observed between the two models indicating comparable restoration of anterior tibial stability at 4 weeks post-operatively. However, comparison to the contralateral limb revealed a significant increase in the impaired healing model, whereas standard ACLR did not differ significantly from the contralateral side. This suggests greater mechanical instability in the impaired healing model (Supplementary Fig. 3).

Having validated the impaired post-ACLR healing, we next examined the bone tunnel, where the healing occurs by formation of fibrovascular granulation interface tissue between the tendon and the bone, followed by remodeling and bone ingrowth. Analysis of Haematoxylin and eosin stained tissue sections using a previously validated scoring system^50^ revealed abnormal and inferior bone-to-tendon healing in the impaired healing model compared to standard ACLR at both 2 and 4 weeks post-operatively, with more pronounced differences at 2 weeks (Fig. 3f,g). Comparison of each score for the 5 parameters of the Tendon-bone Tunnel Healing Scores between the two models showed that the inferior healing was significantly associated with increased fibrous tissues at 2 weeks post-operatively and lower bone-to-tendon integration (head-to-head connection) at 4 weeks post-operatively (Fig. 3h)^50^.

New bone formation within the bone tunnels serves as another quantitative measure of ACLR healing^48^. This was assessed by μCT by measuring the amount of newly formed bone within the tunnels, using a cylindrical volume of interest matching the original tunnel size created with either a 23 G or 21 G needle. μCT analysis of the tibial tunnel in standard ACLR showed progressive new bone formation measured by bone volume fraction, increasing until 4 weeks post-operatively and then plateauing by 8 weeks post-operatively (Fig. 3i). Therefore, we focused our comparative analysis at 4 weeks post-operatively. Although no significant differences in bone volume fraction were observed between the two models, the impaired healing model exhibited increased trabecular number and a trend toward decreased tissue mineral density (Fig. 3j, k). These findings suggest that despite comparable levels of new bone formation, bone-to-tendon integration is compromised in the impaired healing model.

### Increased host-derived immune cells in the impaired healing model

We next investigated the cells and their origins associated with the inferior healing observed in the impaired healing model. Using the MIP-GFP tendon graft/UBC-GFP host transplant model where all green fluorescent cells originate from the host (Fig. 4a), we observed increased host-derived GFP+ cells within the bone tunnels in the impaired healing model compared to standard ACLR (Fig. 4b, c), which was confirmed by image quantification (Fig. 4d). Flow cytometry analysis of tissues harvested from the bone tunnels showed a trend of increased total number of cells in the impaired healing model compared to standard ACLR (Fig. 4f). Notably, there was a significant increase in immune cells, whereas the number of stromal cells remained unchanged (Fig. 4e, f). Moreover, almost all immune cells were derived from the host (Supplementary Fig. 4).

**Figure 4.**
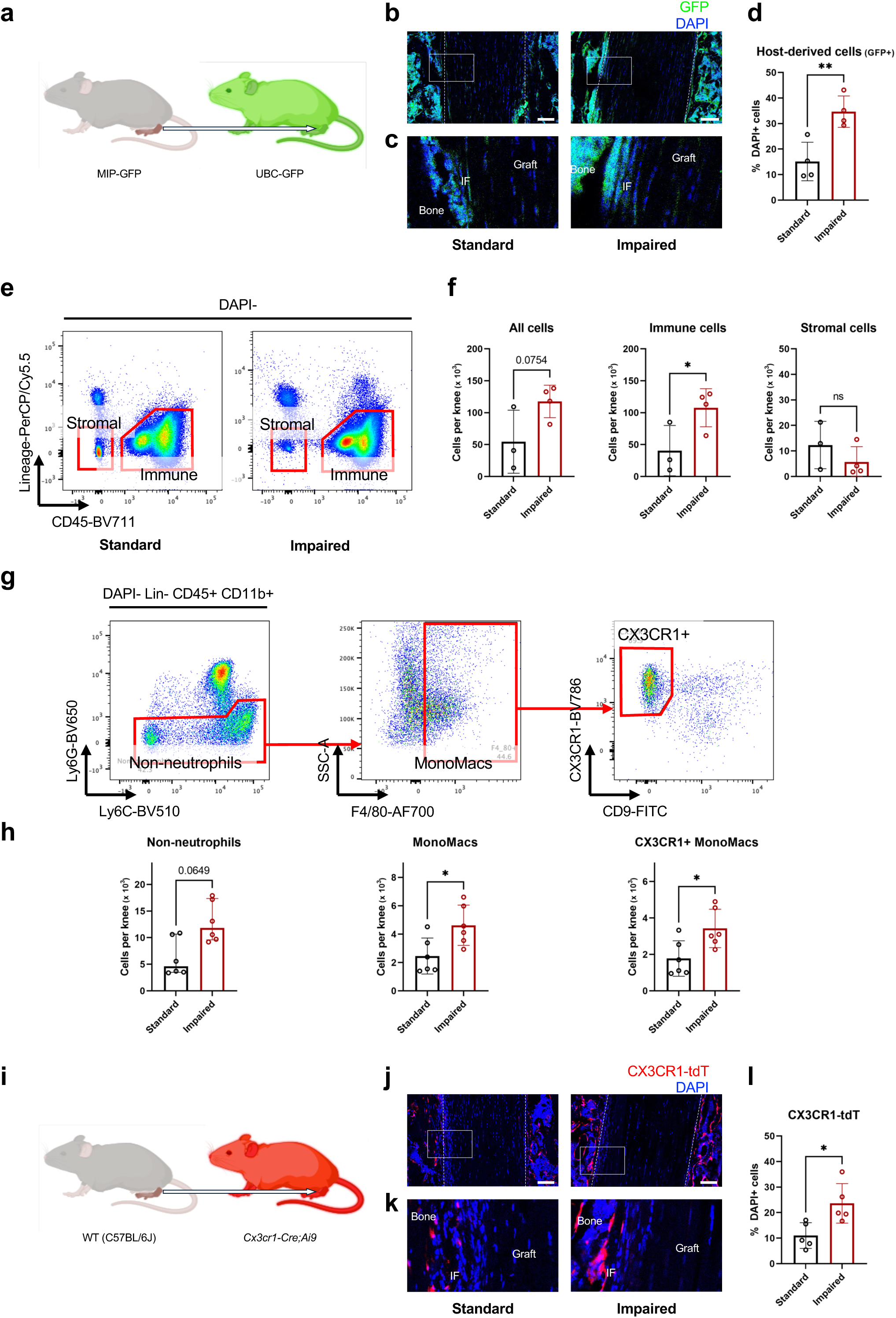
Increased host-derived immune cells in the impaired healing model. **a**, Schematic representation of the MIP-GFP tendon graft/UBC-GFP host transplant model. **b**, Representative immunofluorescence images from 4 replicates using MIP-GFP tendon graft/UBC-GFP host transplant model at 2 weeks post-operatively. Dotted lines indicate the bone tunnel. Scale bar, 100 μm. **c**, Enlarged view of the outlined region (white box) in **b**. **d**, Quantification of immunofluorescent cells (n = 4 per group) show increased host-derived cells (GFP+ cells) within the bone tunnel in the impaired healing model (impaired) compared to standard ACLR. **e**, Representative flow cytometric plots from 3 and 4 replicates for standard ACLR and the impaired healing model, respectively at 2 weeks post-operatively indicating predominance of immune cells in the impaired healing model. **f**, Flow cytometric quantification showing increased immune cells and no changes in lineage (Ter119, CD31, CD3, B220, NK1.1)- CD45-stromal cells in the impaired healing model (n = 4) compared to standard ACLR (n = 3). **g**, Representative flow cytometric plots from 6 replicates and gating strategy at 2 weeks post-operatively for myeloid cells. **h**, Flow cytometric quantification (n = 6 per group) showing increased MonoMacs and CX3CR1+ MonoMacs in the impaired healing model compared to standard ACLR. Each dot represents cells from 3 mice were isolated from the bone tunnels and pooled one sample (**f**, **h**). **i**, Schematic representation of the WT tendon graft/*Cx3cr1-Cre;Ai9* host transplant model. **j**, Representative fluorescence images from 5 replicates using WT tendon graft/*Cx3cr1-Cre;Ai9* host transplant model at 2 weeks post-operatively. Dotted lines indicate the bone tunnel. Scale bar, 100 μm. **k**, Enlarged view of the outlined region (white box) in **j**. **l**, Quantification of fluorescence (n = 5 per group) show increased CX3CR1-tdTomato positive cells within the bone tunnel in the impaired healing model compared to standard ACLR. Bars in the graphs represent the mean and standard deviation except for non-neutrophils where bars represent the median and interquartile range (**h**). \**p* < 0.05; \*\**p* < 0.01; ns, no significance by unpaired *t*-test or Mann-Whitney *U*-test. IF, tendon-bone interface; Graft, tendon graft; DAPI, 4’,6-diamidino-2-phenylindole; SSC, side scatter.

Given our previous findings highlighting the critical role of macrophages in ACLR healing^34, 35^, we next assessed whether there were any differences in the myeloid cells, specifically monocytes and macrophages between the impaired healing model and standard ACLR. As monocytes assume a trajectory of differentiation into macrophages after entering inflamed tissues, we have adopted a ‘MonoMac’ terminology in cases where it is difficult to definitively distinguish these cell types. Among myeloid immune cells, defined as lineage (Ter119, CD31, CD3, B220, NK1.1)-CD45+ CD11b+, both neutrophil and non-neutrophil populations showed a trend toward increased cell number in the impaired healing model. Strikingly, there was a significant increase in the non-neutrophil F4/80+ MonoMac population (Fig. 4h, middle panel). The F4/80+ cells separated into CX3CR1+ CD9-cells (monocytes with CX3CR1-hi cells likely corresponding to non-classical monocytes) and CD9+ cells (likely macrophages) (Fig. 4g, right panel). Notably, the CX3CR1+ CD9-cells were significantly elevated in the impaired healing model (Fig. 4h, right panel).

To corroborate this finding, we employed another model in which a tendon graft from a WT mouse was transplanted into a *CX3CR1-Cre;Ai9* host (Fig. 4i). In this lineage tracing approach, host CX3CR1+ cells express tdTomato and will fluoresce red. Histological analysis revealed increased CX3CR1+ cells within the bone tunnels, especially at the tendon-bone interface, in the impaired healing model compared to standard ACLR (Fig. 4j, k). Image quantification confirmed a significant increase in host-derived CX3CR1+ cells in the impaired healing model at 2 weeks post-operatively (Fig. 4l). Collectively, these results show increased infiltration of CX3CR1+ MonoMacs, which we have previously shown to have an inflammatory phenotype^35^, under conditions of impaired healing.

### scRNA-seq identifies MonoMac clusters in the bone tunnel healing tissues after ACLR

We performed single cell RNA-sequencing (scRNA-seq) to define MonoMac subsets associated with impaired healing and to gain insight into how these cells may compromise the healing process. We harvested tissues from the bone tunnels at 2 weeks post-operatively; cells were pooled from 22 mice in the standard ACLR group, 15 mice in the impaired healing group, and 3 mice in the sham arthrotomy surgery group. Live, non-erythroid, non-lymphoid, non-vascular immune cells (DAPI-Ter119-B220-CD3-NK1.1-CD31-CD45+) were flow-sorted for transcriptomic analysis and a total of 31,288 cells were sequenced using the 10x Genomics platform. After correcting for batch effects across datasets, we applied Uniform Manifold Approximation and Projection (UMAP) for nonlinear dimensionality reduction and conducted graph-based clustering of pooled cells. MonoMacs segregated into 6 distinct clusters (Fig. 5a), which could be classified based upon expression of chemokine receptors CCR2 and CX3CR1, Ly6C, and MHC class II into classical Ly6C-hi monocytes (C1 and C4), non-classical CX3CR1-hi monocytes (C7), MHC class II-hi MonoMacs (C2 and C3; C3 expresses the highest levels of interferon-stimulated genes), and macrophages that expressed *Adgre1* (F4/80), *Trem2* and *C1q* (Fig. 5b-d). Most clusters expressed inflammatory and interferon-stimulated genes (ISGs) such as *Il1b* and *Ifitm3*, in line with previous work showing their inflammatory phenotype^35^. Surprisingly, most clusters expressed *Tgfb1*, which is linked with fibrosis and compromised wound healing^51–53^. *Tgfb1* was most highly expressed in C7 non-classical monocytes, which also expressed the TGF-β receptor *Tgfbr1*. It was also noteworthy that C5 macrophages showed expression of *Spp1*, a canonical marker of pro-fibrotic pathological macrophages^54–57^. These results reveal a profibrotic myeloid cell phenotype that was not clearly apparent in our previous data set that did not include the impaired healing model^35^.

**Figure 5.**
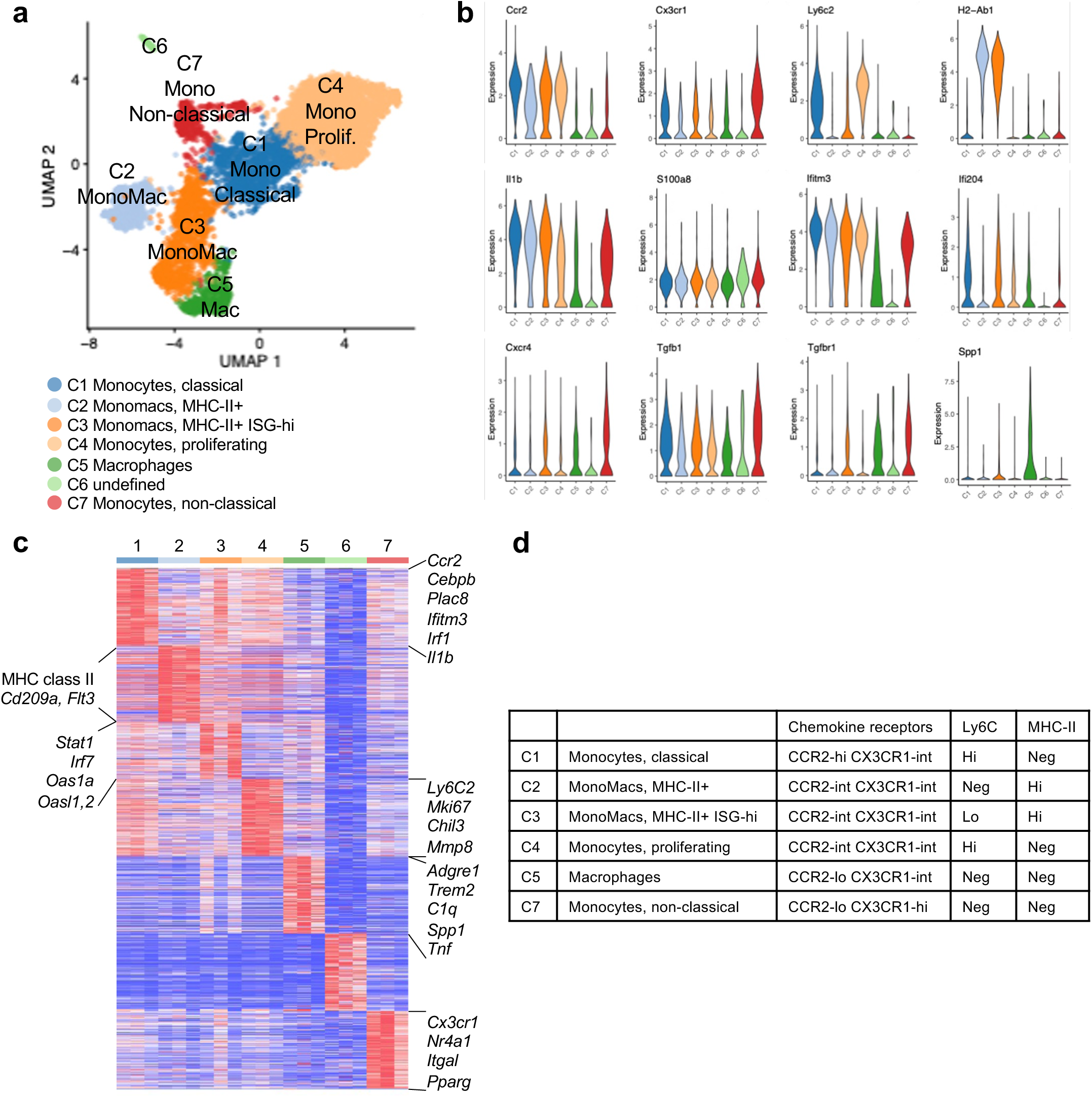
Single-cell RNA-seq identifies MonoMac clusters in the bone tunnel healing tissues after ACLR. Analysis of single cell RNA-sequencing (scRNA-seq) data at 2 weeks post-operatively on cells obtained from 22 mice (standard ACLR model), 15 mice (impaired healing model), and 3 mice (sham arthrotomy surgery). **a**, UMAP projection of scRNA-seq data sub-clustering on MonoMacs. **b**, Violin plots showing expression of the indicated genes. **c**, Heat map showing top genes enriched in the indicated MonoMac clusters. **d,** Table classifying MonoMac subsets based upon expression of chemokine receptors CCR2 and CX3CR1, classical monocyte marker Ly6C and MHC class II (as shown in **b**).

**Figure 6.**
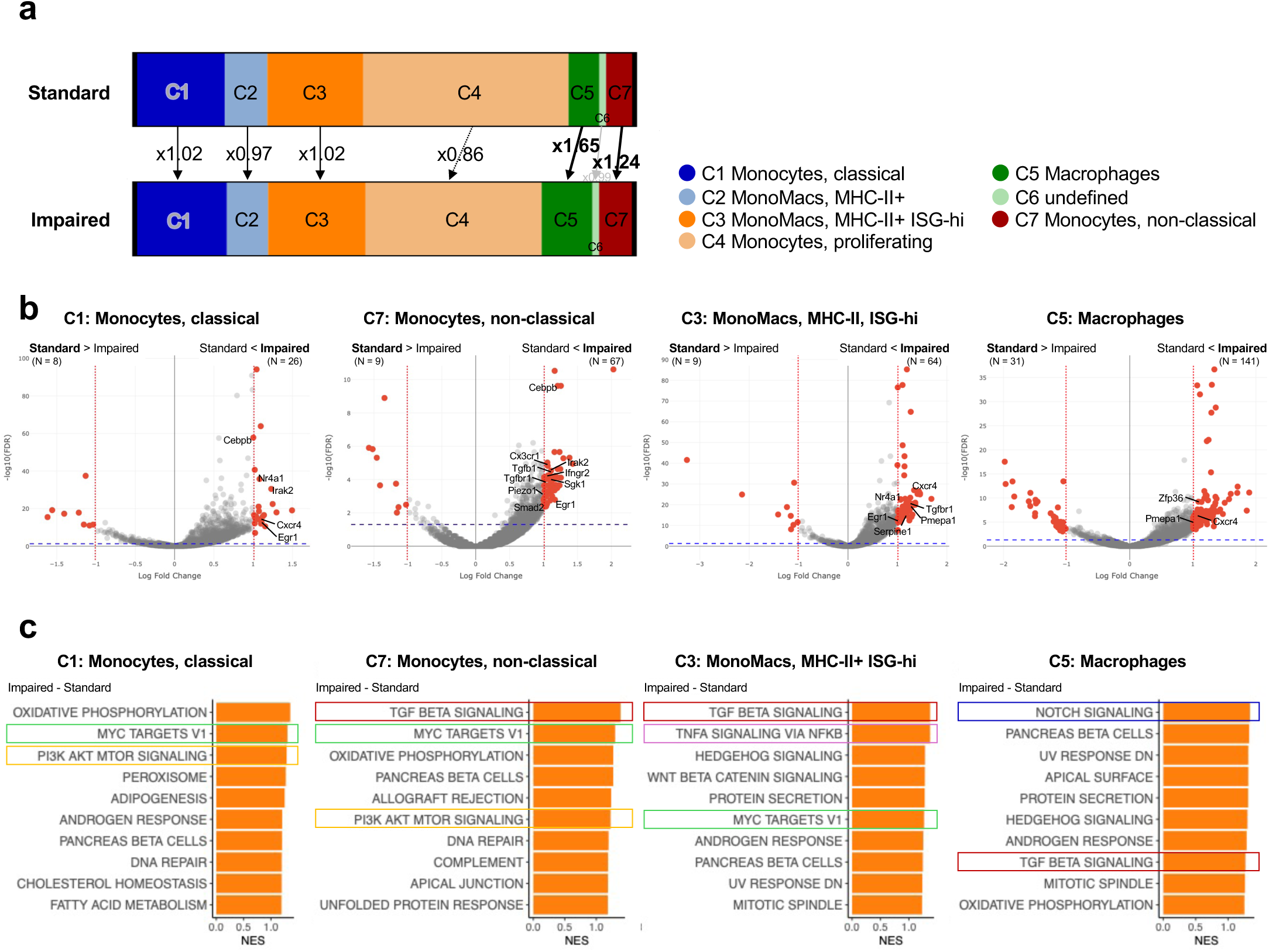
Pathologic activation of MonoMac subsets in the impaired healing model identified by RNA-seq. a, Distribution plot showing relative numbers of MonoMac cell clusters in standard ACLR versus the impaired healing model (impaired). **b**, Volcano plots showing differentially expressed genes between the impaired healing model compared to standard ACLR in the indicated MonoMac clusters. Red dotted line indicates log fold change = 1. Blue dotted line indicates FDR = 0.05. **c**, Pathway analyses using GSEA in the indicated cell clusters.

### Pathologic activation of MonoMac subsets in the impaired healing model identified by RNA-seq

Given the functional importance of CX3CR1+ CCR2+ cells in ACLR healing,^35^ we next assessed differences in the MonoMacs subsets between the standard ACLR and the impaired healing model. At 2 weeks post-operatively, the impaired healing model exhibited a modest increase in non-classical monocytes (C7, 1.24-fold) and a larger increase in macrophages (C5, 1.65-fold) compared to standard ACLR (Fig. 5e). In addition, the pattern of gene expression was significantly altered in all cell subsets in the impaired healing model (Fig. 5f, g). Notably, the non-classical monocytes (C7) expressed elevated levels of fibrotic genes such as *Tgfb1* and *Tgfbr1*, along with genes associated with mechano-transduction pathways, including *Egr1*, *Sgk1* and *Piezo1* (Fig. 5f). These findings are consistent with the design of the impaired healing model, which is predicted to result in a degree of relative micromotion of the graft within the bone tunnels (graft piston or “windshield wiper” effect)^14, 17^, leading to persistent inflammation and fibrovascular granulation tissue formation. Several MonoMac subsets expressed elevated CXCR4, which could potentially mediate monocyte retention and positioning relative to CXCL12-producing stromal cells in tendon-bone interface tissue^58^.

Additionally, in classical and non-classical monocytes (C1 and C7, respectively), the increased expression of *Cebpb* in the impaired healing model was notable as its expression is implicated in the transition of classical monocytes to non-classical monocytes. In classical monocytes (C1) and MHC class II-hi ISG-hi MonoMacs (C3), we observed increased expression of *Nr4a1*, which is also known to be upregulated during monocyte conversion, further supporting phenotypic transition within this compartment. In MHC class II-hi ISG-hi MonoMacs (C3) and macrophages (C5), the impaired healing model showed elevated expression of *Pmepa1*, a regulator of the TGF-β signaling.

In line with the pattern of gene expression described above, gene set enrichment analysis (GSEA) of MonoMac clusters was notable for significant enrichment of TGF-β pathways genes across multiple cell clusters (Fig. 5g). Additional pathways that were commonly enriched in across clusters were PI3K-AKT-mTOR and Myc pathways; MHC class II-hi ISG-hi MonoMacs (C3) also showed elevated TNF-NF-κB pathway activity; and macrophages (C5) showed elevated Notch pathway activity in the impaired healing model relative to standard ACLR. Collectively, these results support the emergence of pro-fibrotic myeloid cell phenotypes associated with impaired post-ACLR healing, and suggest potential roles for mTOR, Myc, Notch and superactivation of NF-kB pathways in pathogenesis.

## DISCUSSION

Our study demonstrates that the early phase of ACLR healing is primarily associated with host-derived cells, mainly immune cells, with tendon graft-derived stromal cells also contributing at the healing tendon-bone interface. Using a clinically relevant murine model of impaired post-ACLR healing induced by a surgical approach, we observed increased infiltration of immune cells, and identified a distinct non-classical monocyte population marked by the expression of CX3CR1 and enriched for fibrotic and mechano-transduction-related gene signatures. Single-cell transcriptomic analysis revealed dynamic transitions among monocyte and macrophage subsets, characterized by a shift toward a pro-fibrotic phenotype in the impaired healing model. Collectively, these findings highlight pathogenic gene expression in host-derived CX3CR1+ non-classical monocytes and additional myeloid cell subsets in impaired post-ACLR healing and suggest these cells compromise the tissue healing process. Thus, targeting their activity may represent a promising therapeutic strategy.

Using a murine ACLR transplant model, we showed that the cells in the bone tunnel healing tissues are host cells likely derived from bone marrow. This finding is consistent with previous transplant ACLR studies in larger animals such as rabbits and pigs^18, 19^. We further identified that the host-derived cells are predominantly immune cells during the early healing phase at 2 weeks post-operatively. Meanwhile, the fate of tendon graft-derived cells has not been directly examined, and their limited role has been generally accepted based on clinical studies reporting similar outcomes between cellular autografts and decellularized allografts^59–64^. In our murine ACLR model, we observed that tendon-derived cells are maintained at the tendon-bone interface, particularly during the early phase of healing, suggesting that they may play a contributory role. This aligns with our concurrent study using a murine ACLR model with decellularized tendon grafts, which also supports a positive role for tendon graft-derived cells in the healing process (manuscript submitted for publication).

To investigate the cellular mechanisms that either promote or dampen ACLR healing, we developed a model of impaired post-ACLR healing using a surgical approach based on the observation that surgical technique is a major contributor of ACLR^9^. Clinical studies have reported higher failure rates with hamstring grafts of smaller diameters^10–12^. Additionally, in a murine tibial prosthesis model, tibial implant osseointegration was impaired when over-sized drill holes were used^65^. In our model, graft fixation was modified to maintain the tendon graft in a deliberately loose state, while other major surgical variables, such as graft harvesting, intra-articular approach, and fixation methods, were kept consistent^66^. Comparing this model with the standard ACLR enabled us to identify key mechanisms underlying impaired post-ACLR healing.

Using a transplant ACLR model, we confirmed through both histological analysis and flow cytometry that the increased cellularity observed in the impaired healing model, associated with the formation of the fibrovascular granulation tissues, is derived from the host. Notably, the increase was primarily attributable to immune cells rather than stromal cells. Together with our concurrent findings demonstrating impaired healing in ACLR using decellularized tendon grafts, where only host-derived cells are present, and previous work showing a deleterious effect of macrophages in the standard ACLR model^34^, the results suggest that an excessive accumulation of host-derived immune cells may be detrimental to ACLR healing. Based on this, we sought to identify the specific host-derived immune cells contributing to inferior outcomes. We have previously shown that macrophages play a key role in ACLR healing^34^, and that targeting a distinct CCR2+ CX3CR1+ macrophage subset expressing inflammatory and IFN-stimulated genes can alter the healing response^35^. Using scRNA-seq of immune cells isolated from healing tissues within the bone tunnels, we identified six distinct MonoMac clusters: three monocyte clusters including classical and non-classical monocytes, two MHC class II-hi MonoMac clusters, and one macrophage cluster. Classical monocytes (C1) represented the predominant population and overlapped with the CCR2+ CX3CR1+ macrophages previously reported by our group^35^. Notably, we observed an increase in non-classical monocytes (C7) in the impaired healing model. These cells were defined by high *Cx3cr1* expression and retained a monocyte identity, lacking differentiation markers such as *Trem2*. They also exhibited enrichment of the TGF-β signaling pathway, indicative of early reprogramming within a fibrotic tissue environment. Similar activation of profibrotic signaling pathways was observed in other MonoMac populations, including macrophages (C5), which were characterized by the expression of *Spp1* and enrichment of TGF-β and Notch pathways.

Interestingly, our prior analysis of MonoMacs at the same timepoint in the standard ACLR model by flow cytometric sorting and bulk RNA sequencing suggested a wound-healing phenotype based upon expression of growth factor genes such as *Pdgfb* and *Igf1*, and angiogenic genes *Vegfa* and *Angpt1*^35^. Collectively, these findings suggest that the abnormal mechanical environment in the impaired healing model subverts a wound-healing cell type to instead assume a deleterious profibrotic role. These cell populations and their associated pathways thus represent potential therapeutic targets for improving ACLR healing.

This study has several limitations. First, all findings are based on murine models of ACL reconstruction. While these models allow for detailed molecular and cellular analyses, differences in anatomy, biomechanics, and immune responses between mice and humans may limit direct translation. The murine ACLR model does not fully recapitulate the complexity of human surgery or graft integration, which should be considered when interpreting the results. Second, although scRNA-seq identified distinct MonoMac populations associated with healing outcomes, different infiltrating cell populations were not analyzed in detail. Finally, the current study did not include therapeutic intervention experiments, which are necessary to validate the functional roles of identified cell populations and pathways. Future studies incorporating therapeutic interventions, such as local delivery of inhibitors or cell-based modulation, will be essential to define actionable strategies for improving graft integration.

In summary, our study demonstrates that host-derived MonoMac subsets, many of which express CX3CR1, may drive the dampened healing observed in the impaired healing ACLR model. Using a combination of histology, flow cytometry, and scRNA-seq, we identified a shift in immune cell composition with increased accumulation of non-classical monocytes marked by reprogramming and elevated TGF-β signaling within a fibrotic microenvironment. Similar fibrotic signatures were seen in other MonoMac populations, including classical monocytes and macrophages. These findings align with prior work highlighting the pivotal role of MonoMac subsets in ACLR healing and extend this understanding by implicating non-classical monocytes as key cell types influenced by the fibrotic environment before macrophage differentiation. These results underscore the importance of tightly regulated immune responses for successful graft integration and suggest that targeting MonoMac subsets and their associated signaling pathways may offer promising therapeutic avenues to enhance ACLR outcomes.

## MATERIALS AND METHODS

### Animals

All experiments were approved by the Institutional Animal Care and Use Committee of Weill Cornell College of Medicine (protocol 2015-0057) and performed in accordance with the National Institutes of Health Guidelines for Care and Use of Laboratory Animals. C57BL/6J (stock 000664), UBC-GFP (stock 004353), Pep Boy (B6 CD45.1, stock 002014), LepR-Cre (stock 008320), Ai9 (stock 007909), Cx3cr1-Cre (stock 025524), and Cx3cr1-GFP (stock 005582) mice were purchased from The Jackson Laboratory (Bar Harbor, ME). MIP-GFP mice were provided by Xu Yang and Mathias Bostrom (HSS). All mice were maintained on a C57BL/6 background. Mice were housed under standard conditions, with a 12-hour light/dark cycle, room temperature at 20.5 °C to 22 °C, and free access to dry laboratory food and water. All experiments were carried out in 12- to 16-week old male mice with at least three animals per group. Euthanasia was carried out by CO_2_ asphyxiation.

### ACL reconstruction surgery

ACLR surgery was conducted on the right knees using previously described methods^39^. In brief, mice were anaesthetized with isoflurane and buprenorphine, and laid in a supine position. Bupivacaine was administered locally for pain relief. A longitudinal incision was made on the medial side of the lower leg, and the flexor digitorum longus tendon (FDL) was detached at the musculotendinous junction. A second incision on the plantar side of the foot exposed the FDL tendon, which was clamped with a hemostatic clip (GEM1521, Synovis Micro Companies Alliance, Birmingham, AL) and transected. A medial parapatellar arthrotomy of the knee joint was performed, and the ACL was severed. Bone tunnels were drilled in the femur and tibia in an isometric configuration, using either a 23 G needle (diameter 0.64 mm, standard) or a 21 G needle (diameter 0.82 mm, impaired healing). The tendon graft was passed through the bone tunnels and sutured to the tibial shaft after a pre-tension of 0.5 N (standard) or no tension (impaired healing). Mice recovered immediately after surgery, exhibited no obvious signs of pain, and were allowed free movement. Buprenorphine was administered post-operatively for pain management.

### Histological analyses

Dissected knees samples were fixed in 10 % neutral buffered formalin (Forma-Scent® Fixative, Cardinal Health, Dublin, OH), decalcified in 0.5 M ethylenediaminetetraacetic acid (VWR, Radnor, PA), cryoprotected with 30 % sucrose, and embedded in O.C.T. compound (Tissue-Tek®, Sakura Finetek, Torrance, CA). Sagittal sections were prepared using Cryofilm (Section-Lab, Yokohama, Japan)^67^, and used for immunohistochemistry and hematoxylin and eosin staining.

For immunohistochemistry, sections were rehydrated in phosphate buffered saline (PBS) and permeabilized with 0.5 % Triton X-100 (Thermo Scientific, Waltham, MA). GFP samples were blocked with 5 % donkey serum, and incubated with anti-GFP primary antibody (1:2000 dilution; ab13970; Abcam, Cambridge, MA) followed by FITC secondary antibody (1:50 dilution; SA1-72000; Invitrogen, Waltham, MA) prior to applying DAPI. Images were acquired using a confocal microscope (Zeiss LSM 880 with Airyscan high-resolution detector, Oberkochen, Germany). Imaging analysis was performed using an automated algorithm on QuPath (0.5.1-x64)^68^. Regions of interest were drawn for the whole bone tunnel or for a width of 100 μm when assessing the tendon-bone interface, and assessed for at least two sections per sample.

Hematoxylin and eosin-stained sections were examined under brightfield microscopy, and representative images were captured with a slide scanner at 20x magnification (Axioscan 7, Zeiss, Oberkochen, Germany). At least three sections per sample were evaluated in a blinded manner using the Tendon-bone Tunnel Healing Scoring system, with median values used for analysis^50^. In brief, the scoring system assessed 5 parameters: graft degeneration, graft remodeling, fibrovascular tissue at the tendon-bone interface, collateral connection between the bone and tendon, and head-to-head bone-to-tendon connection. Each parameter, except for fibrovascular tissue, was scored from 0 to 4, while fibrovascular tissue was scored from 0 to 5, with higher scores indicating better healing.

### Cell isolation

Tissues within the femoral and tibial bone tunnels were harvested using 22 G and 20 G needles for standard ACLR and the impaired healing model, respectively. This harvesting technique captures the fibrovascular healing tissues within the bone tunnels including the tendon graft and a sliver of underlying bone, reflecting the cellular composition of the tendon graft, tendon-bone healing interface, and the reactive newly formed bone^35^. The proximal tibia was harvested for the sham arthrotomy group. Harvested tissues were digested with Collagenase P (1 mg/ml; Roche, Mannheim, Germany), Dispase (2 mg/ml; Gibco, Waltham, MA), and DNase I (0.02 mg/ml; Roche, Mannheim, Germany) for a total of 40 minutes and filtered through a cell strainer. Erythrocytes were removed with ACK lysis buffer (Gibco, Waltham, MA).

### Flow cytometry staining and sorting

Isolated cells were incubated with anti-mouse CD16/32 antibody (TruStain FcX^TM^, Biolegend, San Diego, CA) for 10 minutes to block Fc receptors, followed by staining with primary antibodies including anti-mouse Ter119, B220, CD3, NK1.1, CD31, CD45, CD45.2, CD11b, Ly6G, Ly6C, F4/80, CX3CR1 and CD9 (1:100 dilution) for 15 minutes. Fresh stained cells were run on BD FACSymphony^TM^ A3 Cell Analyzer (BD Biosciences, Franklin Lakes, NJ) and analysis was performed using FlowJo software (v.10.10.0, Tree Star, Ashland, OR), using our previously validated gating strategy^35^.

For cell sorting, isolated cells were first incubated with CD31 and CD45 microbeads (130-097-418 and 130-110-618, respectively; Miltenyi Biotec, Gaithersburg, MD) for 10 minutes each and selected for CD31-CD45+ cells using LS Columns (Miltenyi Biotec). Then, the cells were stained in a similar manner as described above and sorted for Ter119-B220-CD3-NK1.1-CD31-CD45+ on BD FACSymphony^TM^ S6 Cell Sorter (BD Biosciences).

### Biomechanical analysis

Biomechanical testing was conducted as previously described^39^. In brief, operated limbs were stored at -20°C after euthanasia, then thawed at room temperature and dissected to isolate the reconstructed femur-ACL-tibia construct. Samples were potted in microtubes using a filler (Bondo®, 3M, St. Paul, MN) and subjected to tensile loading at a rate of 10 mm/min using a custom-made testing machine. The load to failure (N) for each sample was recorded. Samples were kept moist with saline spray throughout the preparation and testing.

### Gait analysis

Gait analysis was assessed using Digigait^TM^ Imaging System (Mouse Specifics, Framingham, MA), with mice placed on a treadmill at a constant speed (15 cm/s) for a minimum of 10 steps. Stride, stride length and stride frequency were analyzed as representative parameters of gait using the Digigait^TM^ Analysis Software.

### Radiographic analyses

Plain radiographs were taken with the Faxitron UltraFocus DXA (Faxitron Bioptics, Tucson, AZ). Mice were anaesthetized with isoflurane and laid in a supine position to capture both anteroposterior and lateral views. The image settings followed the manufacturer’s recommendations (exposure time 3 – 5.3 seconds at 38 - 45 kV). Osteoarthritis development was evaluated using anteroposterior view radiographs by Kellgren-Lawrence grading scale by in a blinded manner.

Bone formation within the bone tunnel was assessed using a Scanco micro-CT-45 (Scanco Medical, Brüttisellen, Switzerland) as previously described^35^. In brief, knees were dissected, soft tissues removed, and fixed in 10 % neutral buffered formalin before scanning. μCT images were taken with a voxel size of 6 μm (55 kVp, 145 μA, 400 ms integration time). A cylindrical volume of interest, corresponding to the bone tunnel initially created with either 23 G or 21 G needles (diameter 0.64 mm or 0.82 mm, respectively), was identified, and the newly formed bone was measured.

### Single cell RNA sequencing

Cells pooled from 22 mice in standard ACLR, 15 mice in the impaired healing model, and 3 mice in sham arthrotomy surgery were sorted for Ter119-B220-CD3-NK1.1-CD31-CD45-populations. scRNA-seq of a total of 31,288 cells was performed using the 10x Genomics Chromium Single Cell 3’ GEM Library Kit (10x Genomics, Pleasanton, CA), according to the manufacturer’s protocol. Data processing and analysis were conducted using the *scater and scran* packages. Log-normalization and selection of highly variable genes were carried out based on both mean-variance modeling and deviance-based methods. Batch effects were corrected using *Harmony*, and dimensionality reduction was performed using UMAP on Harmony-corrected components. Differentially expressed genes (DEGs) across cell types or clusters were identified using the *scran* package, with significance defined as fold change >1.5 and a Benjamini-Hochberg adjusted p-value <0.05. Enriched pathways within each DEG set were identified using *ClusterProfiler* with Reactome and Gene Ontology databases.

### Statistical Analysis

Data sets were tested for normal distribution using the Shapiro-Wilk normality test. Parametric tests were applied when the data showed normal distribution, while non-parametric tests were used when the data were not normally distributed. Statistical significance was set at p < 0.05. All statistical analyses were performed using Prism 9 Version 9.1.0 (GraphPad Software, Boston, MA).

## Acknowledgements

We thank the micro-CT Core at HSS, Flow Cytometry Core, Genomics Resources Core, and Optical Microscopy Core at Weill Cornell Medicine for their technical support.

## Data and materials availability

The scRNA-seq dataset that was generated by the authors as part of this study has been deposited in the Gene Expression Omnibus database with the accession code GSE303743.

## Notes

### Competing Interest Statement

S.A.R. has received consulting fees from Teladoc, Inc., Enovis-DJO, and Novartis Pharmaceuticals. There are no potential conflicts of interests declared from other authors.

## REFERENCES

1. Kaeding CC, Leger-St-Jean B, Magnussen RA. Epidemiology and Diagnosis of Anterior Cruciate Ligament Injuries. Clin Sports Med 36, 1–8 (2017).

2. Musahl V, Karlsson J. Anterior Cruciate Ligament Tear. N Engl J Med 380, 2341–2348 (2019).

3. Murray MM. Current status and potential of primary ACL repair. Clin Sports Med 28, 51–61 (2009).

4. Scheffler SU, Unterhauser FN, Weiler A. Graft remodeling and ligamentization after cruciate ligament reconstruction. Knee Surg Sports Traumatol Arthrosc 16, 834–842 (2008).

5. Pauzenberger L, Syre S, Schurz M. "Ligamentization" in hamstring tendon grafts after anterior cruciate ligament reconstruction: a systematic review of the literature and a glimpse into the future. Arthroscopy 29, 1712–1721 (2013).

6. Janssen RP, Scheffler SU. Intra-articular remodelling of hamstring tendon grafts after anterior cruciate ligament reconstruction. Knee Surg Sports Traumatol Arthrosc 22, 2102–2108 (2014).

7. Claes S, Verdonk P, Forsyth R, Bellemans J. The "ligamentization" process in anterior cruciate ligament reconstruction: what happens to the human graft? A systematic review of the literature. Am J Sports Med 39, 2476–2483 (2011).

8. Crawford SN, Waterman BR, Lubowitz JH. Long-term failure of anterior cruciate ligament reconstruction. Arthroscopy 29, 1566–1571 (2013).

9. Di Benedetto P, Di Benedetto E, Fiocchi A, Beltrame A, Causero A. Causes of Failure of Anterior Cruciate Ligament Reconstruction and Revision Surgical Strategies. Knee Surg Relat Res 28, 319–324 (2016).

10. Magnussen RA, Lawrence JT, West RL, Toth AP, Taylor DC, Garrett WE. Graft size and patient age are predictors of early revision after anterior cruciate ligament reconstruction with hamstring autograft. Arthroscopy 28, 526–531 (2012).

11. Mariscalco MW, et al. The influence of hamstring autograft size on patient-reported outcomes and risk of revision after anterior cruciate ligament reconstruction: a Multicenter Orthopaedic Outcomes Network (MOON) Cohort Study. Arthroscopy 29, 1948–1953 (2013).

12. Alkhalaf FNA, Hanna S, Alkhaldi MSH, Alenezi F, Khaja A. Autograft diameter in ACL reconstruction: size does matter. SICOT J 7, 16 (2021).

13. Tsuda E, Fukuda Y, Loh JC, Debski RE, Fu FH, Woo SL. The effect of soft-tissue graft fixation in anterior cruciate ligament reconstruction on graft-tunnel motion under anterior tibial loading. Arthroscopy 18, 960–967 (2002).

14. L’Insalata JC, Klatt B, Fu FH, Harner CD. Tunnel expansion following anterior cruciate ligament reconstruction: a comparison of hamstring and patellar tendon autografts. Knee Surg Sports Traumatol Arthrosc 5, 234–238 (1997).

15. Ajuied A, et al. Anterior cruciate ligament injury and radiologic progression of knee osteoarthritis: a systematic review and meta-analysis. Am J Sports Med 42, 2242–2252 (2014).

16. Lindanger L, et al. Predictors of Osteoarthritis Development at a Median 25 Years After Anterior Cruciate Ligament Reconstruction Using a Patellar Tendon Autograft. Am J Sports Med 50, 1195–1204 (2022).

17. Cinque ME, Dornan GJ, Chahla J, Moatshe G, LaPrade RF. High Rates of Osteoarthritis Develop After Anterior Cruciate Ligament Surgery: An Analysis of 4108 Patients. Am J Sports Med 46, 2011–2019 (2018).

18. Bachy M, Sherifi I, Zadegan F, Petite H, Vialle R, Hannouche D. Allograft integration in a rabbit transgenic model for anterior cruciate ligament reconstruction. Orthop Traumatol Surg Res 102, 189–195 (2016).

19. Takeuchi H, et al. Temporal Changes in Cellular Repopulation and Collagen Fibril Remodeling and Regeneration After Allograft Anterior Cruciate Ligament Reconstruction: An Experimental Study Using Kusabira-Orange Transgenic Pigs. Am J Sports Med 44, 2375–2383 (2016).

20. Goertzen MJ, Buitkamp J, Clahsen H, Mollmann M. Cell survival following bone-anterior cruciate ligament-bone allograft transplantation: DNA fingerprints, segregation, and collagen morphological analysis of multiple markers in the canine model. Arch Orthop Trauma Surg 117, 208–214 (1998).

21. Jackson DW, Simon TM. Donor cell survival and repopulation after intraarticular transplantation of tendon and ligament allografts. Microsc Res Tech 58, 25–33 (2002).

22. Jackson DW, Simon TM, Kurzweil PR, Rosen MA. Survival of cells after intra-articular transplantation of fresh allografts of the patellar and anterior cruciate ligaments. DNA-probe analysis in a goat model. J Bone Joint Surg Am 74, 112–118 (1992).

23. Min BH, Han MS, Woo JI, Park HJ, Park SR. The origin of cells that repopulate patellar tendons used for reconstructing anterior cruciate ligaments in man. J Bone Joint Surg Br 85, 753–757 (2003).

24. Webster DA, Werner FW. Freeze-dried flexor tendons in anterior cruciate ligament reconstruction. Clin Orthop Relat Res, 238–243 (1983).

25. Kanazawa T, Soejima T, Murakami H, Inoue T, Katouda M, Nagata K. An immunohistological study of the integration at the bone-tendon interface after reconstruction of the anterior cruciate ligament in rabbits. J Bone Joint Surg Br 88, 682–687 (2006).

26. Kawamura S, Ying L, Kim HJ, Dynybil C, Rodeo SA. Macrophages accumulate in the early phase of tendon-bone healing. J Orthop Res 23, 1425–1432 (2005).

27. Scranton PE, Jr., Lanzer WL, Ferguson MS, Kirkman TR, Pflaster DS. Mechanisms of anterior cruciate ligament neovascularization and ligamentization. Arthroscopy 14, 702–716 (1998).

28. Weiler A, Unterhauser FN, Bail HJ, Huning M, Haas NP. Alpha-smooth muscle actin is expressed by fibroblastic cells of the ovine anterior cruciate ligament and its free tendon graft during remodeling. J Orthop Res 20, 310–317 (2002).

29. Meller R, et al. Graft remodeling during growth following anterior cruciate ligament reconstruction in skeletally immature sheep. Arch Orthop Trauma Surg 129, 1037–1046 (2009).

30. Unterhauser FN, Bail HJ, Hoher J, Haas NP, Weiler A. Endoligamentous revascularization of an anterior cruciate ligament graft. Clin Orthop Relat Res, 276–288 (2003).

31. Yoshikawa T, Tohyama H, Enomoto H, Matsumoto H, Toyama Y, Yasuda K. Expression of vascular endothelial growth factor and angiogenesis in patellar tendon grafts in the early phase after anterior cruciate ligament reconstruction. Knee Surg Sports Traumatol Arthrosc 14, 804–810 (2006).

32. Kamalitdinov TB, et al. Amplifying Bone Marrow Progenitors Expressing alpha-Smooth Muscle Actin Produce Zonal Insertion Sites During Tendon-to-Bone Repair. J Orthop Res 38, 105–116 (2020).

33. Hagiwara Y, Dyrna F, Kuntz AF, Adams DJ, Dyment NA. Cells from a GDF5 origin produce zonal tendon-to-bone attachments following anterior cruciate ligament reconstruction. Ann N Y Acad Sci 1460, 57–67 (2020).

34. Hays PL, et al. The role of macrophages in early healing of a tendon graft in a bone tunnel. J Bone Joint Surg Am 90, 565–579 (2008).

35. Fujii T, et al. Distinct Inflammatory Macrophage Populations Sequentially Infiltrate Bone-to-Tendon Interface Tissue After Anterior Cruciate Ligament (ACL) Reconstruction Surgery in Mice. JBMR Plus 6, e10635 (2022).

36. Wada S, et al. Remodeling Process of the Tendon Graft After Anterior Cruciate Ligament Reconstruction: Comprehensive Analysis With RNA Sequencing in a Murine Model. J Orthop Res 43, 1122–1131 (2025).

37. Nakagawa Y, et al. Duration of postoperative immobilization affects MMP activity at the healing graft-bone interface: Evaluation in a mouse ACL reconstruction model. J Orthop Res 37, 325–334 (2019).

38. Deng XH, et al. Expression of Signaling Molecules Involved in Embryonic Development of the Insertion Site Is Inadequate for Reformation of the Native Enthesis: Evaluation in a Novel Murine ACL Reconstruction Model. J Bone Joint Surg Am 100, e102 (2018).

39. Camp CL, et al. Timing of Postoperative Mechanical Loading Affects Healing Following Anterior Cruciate Ligament Reconstruction: Analysis in a Murine Model. J Bone Joint Surg Am 99, 1382–1391 (2017).

40. Dagher E, Hays PL, Kawamura S, Godin J, Deng XH, Rodeo SA. Immobilization modulates macrophage accumulation in tendon-bone healing. Clin Orthop Relat Res 467, 281–287 (2009).

41. Gulotta LV, Kovacevic D, Ying L, Ehteshami JR, Montgomery S, Rodeo SA. Augmentation of tendon-to-bone healing with a magnesium-based bone adhesive. Am J Sports Med 36, 1290–1297 (2008).

42. Gulotta LV, Rodeo SA. Biology of autograft and allograft healing in anterior cruciate ligament reconstruction. Clin Sports Med 26, 509–524 (2007).

43. Ma CB, et al. Bone morphogenetic proteins-signaling plays a role in tendon-to-bone healing: a study of rhBMP-2 and noggin. Am J Sports Med 35, 597–604 (2007).

44. Rodeo SA, et al. The effect of osteoclastic activity on tendon-to-bone healing: an experimental study in rabbits. J Bone Joint Surg Am 89, 2250–2259 (2007).

45. Anderson K, Seneviratne AM, Izawa K, Atkinson BL, Potter HG, Rodeo SA. Augmentation of tendon healing in an intraarticular bone tunnel with use of a bone growth factor. Am J Sports Med 29, 689–698 (2001).

46. Rodeo SA, Kawamura S, Kim HJ, Dynybil C, Ying L. Tendon healing in a bone tunnel differs at the tunnel entrance versus the tunnel exit: an effect of graft-tunnel motion? Am J Sports Med 34, 1790–1800 (2006).

47. Kleiner JB, Amiel D, Roux RD, Akeson WH. Origin of replacement cells for the anterior cruciate ligament autograft. J Orthop Res 4, 466–474 (1986).

48. Fu SC, Cheuk YC, Yung SH, Rolf CG, Chan KM. Systematic Review of Biological Modulation of Healing in Anterior Cruciate Ligament Reconstruction. Orthop J Sports Med 2, 2325967114526687 (2014).

49. Atarod M, Frank CB, Shrive NG. Increased meniscal loading after anterior cruciate ligament transection in vivo: a longitudinal study in sheep. Knee 22, 11–17 (2015).

50. Lui PP, Ho G, Lee YW, Ho PY, Lo WN, Lo CK. Validation of a histologic scoring system for the examination of quality of tendon graft to bone tunnel healing in anterior cruciate ligament reconstruction. Anal Quant Cytol Histol 33, 36–49 (2011).

51. Meng XM, Nikolic-Paterson DJ, Lan HY. TGF-beta: the master regulator of fibrosis. Nat Rev Nephrol 12, 325–338 (2016).

52. Irma J, Kartasasmita AS, Kartiwa A, Irfani I, Rizki SA, Onasis S. From Growth Factors to Structure: PDGF and TGF-beta in Granulation Tissue Formation. A Literature Review. J Cell Mol Med 29, e70374 (2025).

53. Liarte S, Bernabe-Garcia A, Nicolas FJ. Role of TGF-beta in Skin Chronic Wounds: A Keratinocyte Perspective. Cells 9, (2020).

54. Morse C, et al. Proliferating SPP1/MERTK-expressing macrophages in idiopathic pulmonary fibrosis. Eur Respir J 54, (2019).

55. Hoeft K, et al. Platelet-instructed SPP1(+) macrophages drive myofibroblast activation in fibrosis in a CXCL4-dependent manner. Cell Rep 42, 112131 (2023).

56. Jiang Y, et al. Macrophages in organ fibrosis: from pathogenesis to therapeutic targets. Cell Death Discov 10, 487 (2024).

57. Uhlig M, Billig S, Wienhold J, Schumacher D. Pro-Fibrotic Macrophage Subtypes: SPP1+ Macrophages as a Key Player and Therapeutic Target in Cardiac Fibrosis? Cells 14, (2025).

58. Chen H, et al. Pleiotropic Roles of CXCR4 in Wound Repair and Regeneration. Front Immunol 12, 668758 (2021).

59. Hayback G, Raas C, Rosenberger R. Failure rates of common grafts used in ACL reconstructions: a systematic review of studies published in the last decade. Arch Orthop Trauma Surg 142, 3293–3299 (2022).

60. Mascarenhas R, et al. Is there a higher failure rate of allografts compared with autografts in anterior cruciate ligament reconstruction: a systematic review of overlapping meta-analyses. Arthroscopy 31, 364–372 (2015).

61. Cvetanovich GL, et al. Hamstring autograft versus soft-tissue allograft in anterior cruciate ligament reconstruction: a systematic review and meta-analysis of randomized controlled trials. Arthroscopy 30, 1616–1624 (2014).

62. Yao LW, et al. Patellar tendon autograft versus patellar tendon allograft in anterior cruciate ligament reconstruction: a systematic review and meta-analysis. Eur J Orthop Surg Traumatol 25, 355–365 (2015).

63. Mariscalco MW, Magnussen RA, Mehta D, Hewett TE, Flanigan DC, Kaeding CC. Autograft versus nonirradiated allograft tissue for anterior cruciate ligament reconstruction: a systematic review. Am J Sports Med 42, 492–499 (2014).

64. Lamblin CJ, Waterman BR, Lubowitz JH. Anterior cruciate ligament reconstruction with autografts compared with non-irradiated, non-chemically treated allografts. Arthroscopy 29, 1113–1122 (2013).

65. Kuyl EV, et al. Inhibition of PAD4 mediated neutrophil extracellular traps prevents fibrotic osseointegration failure in a tibial implant murine model : an animal study. Bone Joint J 103-B, 135–144 (2021).

66. Hooijmans CR, Leenaars M, Ritskes-Hoitinga M. A gold standard publication checklist to improve the quality of animal studies, to fully integrate the Three Rs, and to make systematic reviews more feasible. Altern Lab Anim 38, 167–182 (2010).

67. Dyment NA, et al. High-Throughput, Multi-Image Cryohistology of Mineralized Tissues. J Vis Exp, (2016).

68. Bankhead P, et al. QuPath: Open source software for digital pathology image analysis. Sci Rep 7, 16878 (2017).

69. Yang R, et al. A single-cell atlas depicting the cellular and molecular features in human anterior cruciate ligamental degeneration: A single cell combined spatial transcriptomics study. Elife 12, (2023).

70. Iwata S, Suda Y, Nagura T, Matsumoto H, Otani T, Toyama Y. Dynamic instability during stair descent in isolated PCL-deficient knees: what affects abnormal posterior translation of the tibia in PCL-deficient knees? Knee Surg Sports Traumatol Arthrosc 15, 705–711 (2007).

