## Supplemental Figures for "Pathologic profibrotic activation of distinct myeloid cell subsets in a model of impaired healing after anterior cruciate ligament reconstruction"

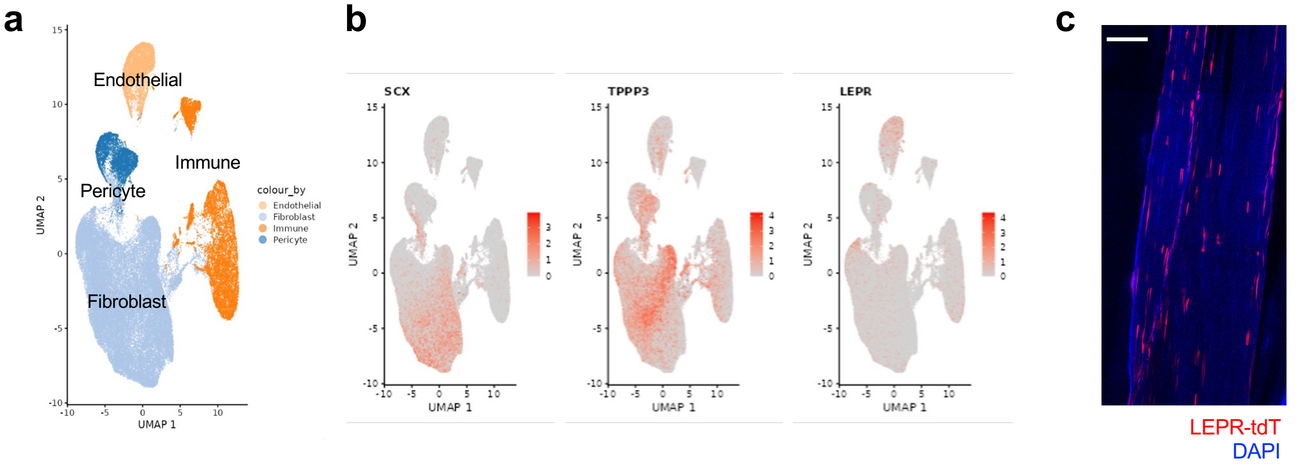


**Supplementary Figure 1. Leptin receptor (LEPR) is expressed in ligament-derived cells including fibroblasts**. **a**, UMAP projection of scRNA-seq data on cells derived from human anterior cruciate ligaments (Genome Sequence Archive for Human, PRJCA014157)[62]. **b**, Feature plots showing well-known marker genes of tendon/ligament-derived cells such as *SCX* and *TPPP3*, and *LEPR*. **c**, Representative fluorescence image from 2 replicates of a flexor digitorum longus tendon from a naïve *Lepr-Cre;Ai9* mouse. Scale bar, 100 μm. DAPI, 4',6-diamidino-2-phenylindole.

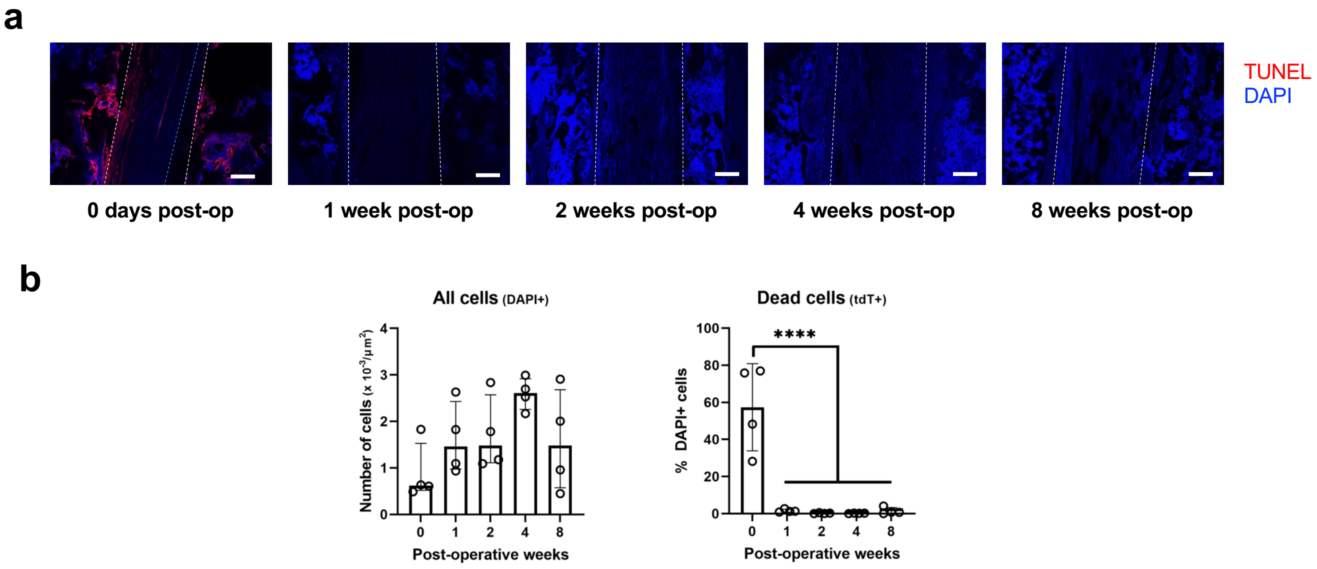


**Supplementary Figure 2. Tendon graft-derived cells are maintained after ACLR surgery. a**, Representative fluorescence images from 4 replicates of TUNEL staining. Dotted white lines indicate the bone tunnel. Dotted blue lines indicate the edge of the tendon graft. Scale bar, 100 μm. **b**, Quantification of fluorescence (n = 4 per timepoint) show no significant changes in the total number of DAPI+ cells within the bone tunnels. TUNEL positive cells are detected immediately after the surgery (0 days pos-op) only. Bars in the graphs represent the median and interquartile range (all cells) or mean and standard deviation (dead cells). ****$p < 0.0001$ by one-way ANOVA with Tukey's post hoc test. TUNEL, terminal deoxynucleotidyl transferase dUTP nick end labeling; DAPI, 4',6-diamidino-2-phenylindole.

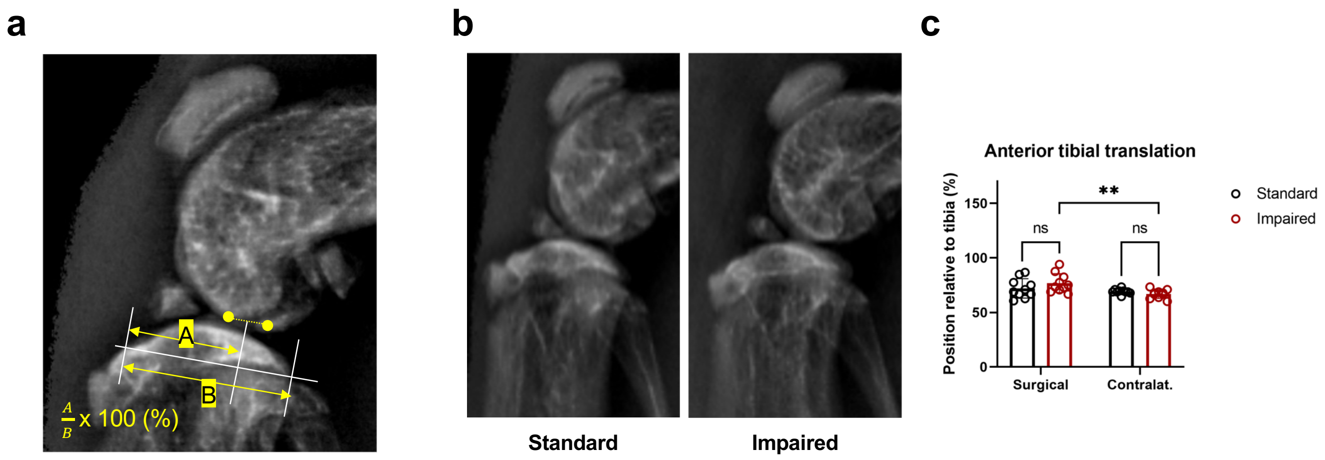


**Supplementary Figure 3. Insufficient restoration of anterior tibial stability in the impaired healing model**. **a**, Midpoint measurement method: the midpoint of the nearest points of the femoral condyles to the tibia were defined. Then the relative distance of the midpoint from the anterior edge (A) to the whole length of the tibial joint surface (B) was calculated as a percentage. **b**, Representative un-stressed lateral view plain radiographs of standard ACLR and the impaired healing model at 4 weeks post-operatively. **c**, Anterior tibial translation show no significant difference between the two models (n = 10 per group). However, it was increased in the impaired healing model compared to the contralateral side (contralat.). Bars in the graph represent the mean and standard deviation. $^{**}p < 0.01$; ns, no significance by two-way ANOVA with Tukey's post hoc test.

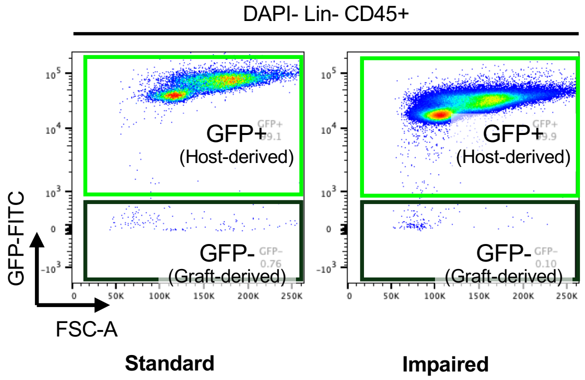


**Supplementary Figure 4. Immune cells are predominantly derived from the host.** Representative flow cytometric plots at 2 weeks post-operatively indicating dominance of GFP+ host-derived cells in immune cells. FSC, forward scatter.

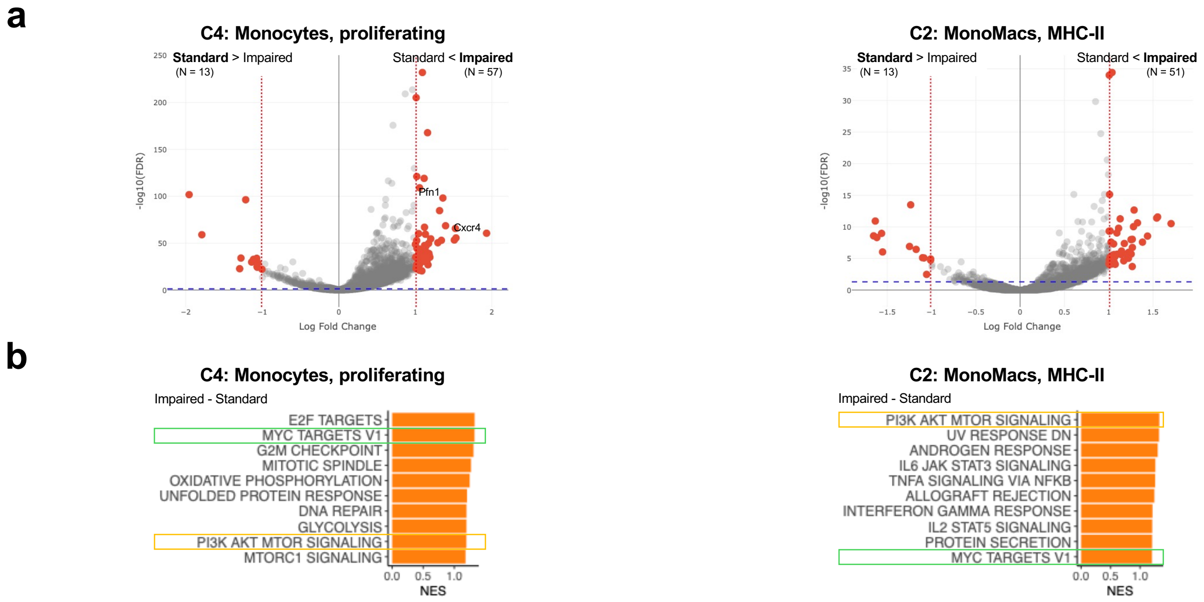


**Supplementary Figure 5. Differentially expressed genes and pathways in proliferating monocytes and MHC class II-hi MonoMacs between standard ACLR and the impaired healing model.** **a**, Volcano plots showing differentially expressed genes between the impaired healing model compared to standard ACLR in proliferating monocytes (C4) and MHC class II-hi MonoMacs (C2). Two-sided *t*-test. Red dotted line indicates log fold change = 1. Blue dotted line indicates FDR = 0.05. **b**, Pathway analyses using GSEA show increased PI3K-AKT-mTOR and Myc signaling in the impaired healing model compared to standard ACLR.
